# Reproductive barriers as a byproduct of gene network evolution

**DOI:** 10.1101/2020.06.12.147322

**Authors:** Chia-Hung Yang, Samuel V. Scarpino

## Abstract

Molecular analyses of closely related taxa have increasingly revealed the importance of higher-order genetic interactions in explaining the observed pattern of reproductive isolation between populations. Indeed, both empirical and theoretical studies have linked the process of speciation to complex genetic interactions. Gene Regulatory Networks (GRNs) capture the inter-dependencies of gene expression and encode information about an individual’s phenotype and development at the molecular level. As a result, GRNs can–in principle–evolve via natural selection and play a role in non-selective, evolutionary forces. Here, we develop a network-based model, termed the pathway framework, that considers GRNs as a functional representation of coding sequences. We then simulated the dynamics of GRNs using a simple model that included natural selection, genetic drift, and sexual reproduction and found that reproductive barriers can develop rapidly between allopatric populations experiencing identical selection pressure. Further, we show that alleles involved in reproductive isolation can predate the allopatric separation of populations and that the number of interacting loci involved in genetic incompatibilities, i.e., the order, is often high simply as a by-product of the networked structure of GRNs. Finally, we discuss how results from the pathway framework are consistent with observed empirical patterns for genes putatively involved in post-zygotic isolation. Taken together, this study adds support for the central role of gene networks in speciation and in evolution more broadly.

## Introduction

Over the past 100 years, the role of reproductive isolation due to genetic incompatibilities has received considerable attention in both the empirical and theoretical literature on speciation (Rieseberg et al., 1996; Coyne and Allen Orr, 1998; Marques et al., 2019; Satokangas et al., 2020). Through this work, it is widely accepted that divergent selection on *de novo* mutations in geographically isolated populations can facilitate speciation, as originally theorized by Bateson (1909); Dobzhansky (1936); Muller (1942). Despite well-established examples from *Drosophila* (Brideau et al., 2006; Wittkopp et al., 2008), *Xiphophorus* (Wittbrodt et al., 1989; Powell et al., 2020), *Oryza* (Yamamoto et al., 2010), *Arabidopsis* (Bikard et al., 2009), and *Mus* (Davies et al., 2016), the genetics and evolutionary history of incompatibilities are typically far more complex and/or less well understood than what is suggested by classical models (Noor and Feder, 2006; Lowry et al., 2008; Presgraves, 2010; Wolf et al., 2010; Nosil and Schluter, 2011; Seehausen et al., 2014; Marques et al., 2019; Dagilis and Matute, 2020).

Post-zygotic, genetic isolation is thought to occur due to epistatic interaction between loci, where alleles arise and fix in allopatry prior to secondary contact, e.g., the Bateson-Dobzhansky-Muller (BDM) model (Bateson, 1909; Dobzhansky, 1936; Muller, 1942). However, many incompatibilities uncovered using high-throughput molecular analyses (Castillo and Barbash, 2017; Kuzmin et al., 2018; Schumer et al., 2018; Vaid and Laitinen, 2019) and quantitative trait locus (QTL) mapping (Moyle and Nakazato, 2008; Turner et al., 2014; Chae et al., 2014; Lowry et al., 2015; Wang et al., 2015), do not conform to the processes assumed by the BDM model. In particular, in both natural populations and model organisms, studies have found that reproductive barriers often exist between allopatric populations experiencing similar selection pressures and that many of the alleles underlying genetic incompatibility predate the allopatric separation of populations (Schluter, 2009; Han et al., 2017; Guerrero and Hahn, 2017; Marques et al., 2019; Jamie and Meier, 2020). Both the lack of divergent selection and the role of standing genetic variation are clear violations of the BDM model. As a result, reconciling theoretical models of how and why genetic incompatibilities arise with emperical data on the molecular genetics of post-zygotic, reproductive isolation is of profound importance (Marques et al., 2019; Satokangas et al., 2020).

Analytical and computational models have proposed theoretical explanations for the observed patterns of complex genetic interaction underlying post-zygotic isolation. A collection of models considers *de-novo* mutations at the population level and the accompanying accumulation of hybrid incompatibilities. For example, Orr (1995) predicted that the number of incompatibilities should increase faster than linearly with the number of substitutions. The study by Orr also suggested higher prevalence of complex genetic interactions than simple pairwise incompatibilities. This so-called “snowballing” effect has been further extended by incorporating protein-protein interaction and RNA folding (Livingstone et al., 2012; Kalirad and Azevedo, 2017). Similarly, Barton (2001) demonstrated that stabilizing selection can generate hybrid incompatibility between allopatric populations using a quantitative genetics models.

The substitution-based approaches, nevertheless, are often at odds with emerging data on the evolutionary history of alleles involved in reproductive isolation (Marques et al., 2019; Satokangas et al., 2020). In addition, many models make an implicit assumption that two allopatric lineages only differ by fixed alleles, which does not capture the empirical diversity among individuals’ gene expression (Kelly et al., 2017; Tyler et al., 2017; Gould et al., 2018; Mogil et al., 2018; Ryu et al., 2019) nor the observed importance of regulatory disruption and standing genetic variation in generating reproductive isolation (Hopkins and Rausher, 2011; Wittkopp and Kalay, 2012; Guerrero et al., 2016; Rougeux et al., 2019; Morgan et al., 2020). More importantly, substitutions originating from *de-novo* mutations fail to explain the recent evidence that alleles underlying reproductive barriers often predate speciation events and can evolve along parallel evolutionary trajectories (Kaeuffer et al., 2012; Sicard et al., 2015; Meier et al., 2017; Nelson and Cresko, 2018; Wang et al., 2019; Duranton et al., 2019; Marques et al., 2019).

Another class of computational approaches focuses on the regulation structure that is potentially responsible for complex genetic interactions and resulting incompatibilities. Specifically, researchers consider the evolution of gene regulatory networks (GRNs), which describe the inter-dependencies between gene expression and encode information about both genotype and phenotype. First, Johnson and Porter (2000) simulated a single linear regulatory pathway as a sequence of matching functions for binding sites, which resulted in reduced hybrid fitness compared to non-epistatic models. Next, Palmer and Feldman (2009) explored the developmental process where the expression of gene products was iteratively determined through the regulatory networks. Their model demonstrated that, largely as a consequence of the diverse set of possible development pathways, hybrid incompatibilities due to disrupted GRNs could evolve rapidly. More recently, Schiffman and Ralph (2018) modeled gene networks as linear control systems and demonstrated that reproductive isolation can be a consequence of parallel evolution of GRNs with equivalent mechanism. Lastly, Blanckaert et al. (2020) showed the importance of higher-order interactions and cryptic epistasis for the evolution of reproductive isolation in the presence of gene flow.

The implications from these GRN models are not mere outcomes of layering complexity onto existing approaches. Instead, GRNs are a natural extension from lower-dimensional models due to their close relationship with coding sequences. Ideally, and hypothetically given “omniscience” over the genomes–including comprehension of every fundamental interaction between molecules–one can reconstruct inter-dependencies among genes and obtain GRNs from a bottom-up approach. Of course, this ambition is far from practical and even sounds like a fantasy. Yet, it shows that GRNs are essentially a direct abstraction of the genome sequence. Furthermore, this abstraction is central to the omnigenic perspective of complex traits (Boyle et al., 2017). GRNs therefore bridge the gap between inheritance factors and physiological traits, whose dynamics over generations then becomes a candidate for understanding the genetics of speciation due to genetic incompatibilities.

To investigate the role of complex genetic interactions in the speciation process, we develop a network-science model for the evolution of GRNs which specifically focuses on the inherited molecular pathways encoded in them. Our approach, termed the pathway framework, considers GRNs as a functional representation of genotype-to-phenotype maps, where proteins are “nodes” in the network and alleles of loci are “edges.” Using this framework, we show how a simple model, which includes sexual reproduction, genetic drift, and natural selection, can drive a rapid increase in reproductive isolation between allopatric populations from standing genetic variation under identical selection pressure. Additionally, we find that genetic incompatibilities can frequently involve many loci, i.e., be of higher order, simply as a by-product of GRN evolution. Finally, we conclude the functional redundancy of GRNs is critical for the rapid emergence of reproductive isolation during population divergence.

## Results

### The pathway framework: networks as a functional representation of genetic interactions

Gene interaction networks are conventionally built such that genes are “nodes” and interactions between genes are “edges” or links, for examples see Tong et al. (2004); Schlitt and Brazma (2007); Langfelder and Horvath (2008). Here, we propose an alternative methodology–termed the *pathway framework*–for constructing gene interaction networks. The key idea of the pathway framework is to conceptualize genes, or alleles of genes, as “black boxes” that encapsulate how their expression is regulated. More precisely, the pathway framework transforms alleles of genes into directed edges pointing from nodes that are activator/repressor molecules, e.g., transcription factors, and nodes that represent gene products, e.g., proteins. In Fig 1 we show how: a.) a gene is activated by a transcription factor and generates a protein product (top-right), b.) two genes interact via a transcription factor created by one gene that activates the other (middle-right), and c.) genes can interact via shared transcription factors (bottom-right). As a result of its flexibility, arbitrarily complex genetic interactions can be encoded as “pathways” through a gene interaction network.

**Figure 1:**
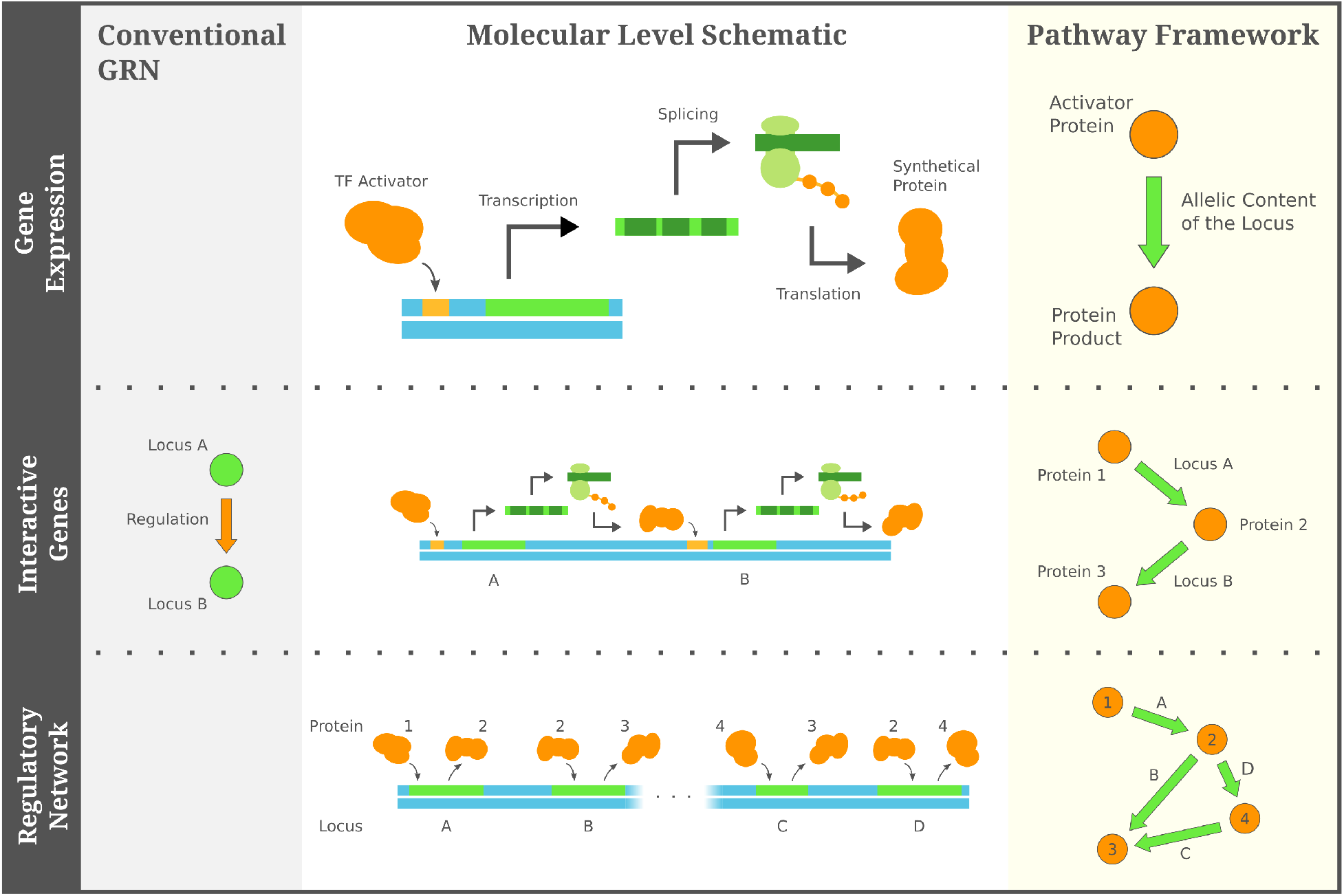
The pathway framework captures complex genetic interactions through consecutive regulatory pathways. In contrast to directly representing genetic interactions, the pathway framework abstracts the regulation and expression of genes as black boxes. If we consider the example of regulation by transcription factors, the pathway framework turns alleles of genes into edges between the transcription factors and the resulting protein products, and regulatory interactions between genes are encapsulated by consecutive pathways through the network.

Importantly, while our proposed representation is closely related to conventional gene interaction networks (and a direct mapping between the two always exists when considering interactions mediated by a single class of molecules, e.g., proteins), the pathway framework is often either a more compact and/or informative representation. For example, anytime a gene is regulated by a protein product from another gene, the conventional framework usually includes redundancy that does not appear in the pathway framework, and the pathway framework will capture information not present in the conventional construction, e.g., see Fig 2. Because the computational complexity of network analyses often scales non-linearly with the number of edges, switching to the pathway framework can facilitate a more robust exploration of model space.

**Figure 2:**
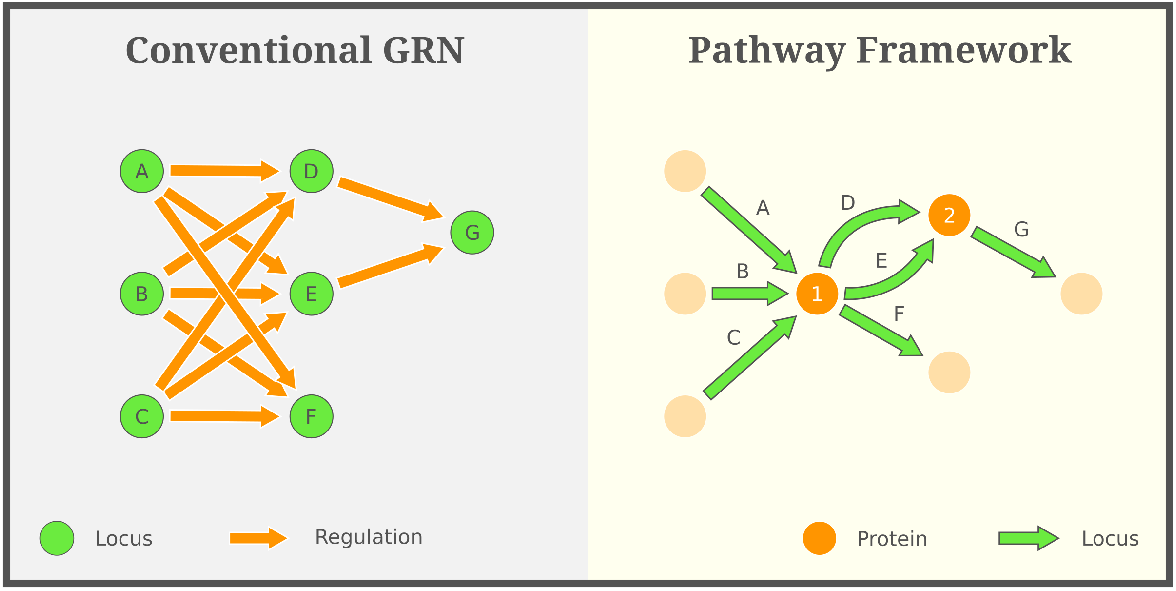
The pathway framework is often a more compact representation. Because the pathway framework directly encodes the expression pattern of genes, it can contain more information than the “conventional” approach to constructing GRNs. When considering genetic interactions that are mediated by a single class of molecules, e.g., one gene being regulated by the protein product of another, the pathway framework takes advantage of this information and presents genetic interactions in a more compact format. Conversely, a conventional GRN lacks the specific regulatory context, and thus it has to present all pairs of interacting genes as individual edges, rather than summarizing these interactions by a smaller set of protein mediators. More technically, the pathway framework and a conventional GRN correspond to the first- and second-order de Bruijn graph (De Bruijn, 1946) respectively, where higher orders usually introduce redundant elements and additional computational complexity.

The pathway framework further highlights how phenotypes are a product of both genetics and the environment (not all nodes in the pathway framework need be gene products). Concentrating on the molecular basis of physiological traits, a phenotype can be thought of as the biochemical status of a universal collection of nodes in the pathway framework, e.g., gene products such as proteins or environmental stimuli. Therefore, under the pathway framework, the development of a phenotype can be viewed as an iterative process of chemical signals propagating through woven pathways built from groups of “inherited metabolisms” and external signals from the environment. As a result, the pathway framework can readily capture genetic, environment, and gene x environment effects in the same network.

### Evolutionary mechanisms under the pathway framework

Although in its most abstract state, the pathway framework can include nodes that are not proteins and also nodes that are not directly involved in gene regulation; here, we focus on the evolution of GRNs where all nodes are proteins directly involved in transcriptional regulation. To model and simulate the evolution of GRNs, this version of the pathway framework translates evolutionary mechanisms–such as mutation, independent assortment, recombination, and gene duplication–into graphical operations on the gene networks. These graphical operations focus on edges in the GRNs, while the underlying node set is held constant because the nodes represent all *possibly existing* proteins in the organism. Because mutation of a locus can potentially alter its protein product and/or the transcription factor binding region(s), we consider mutation as a rewiring process where the incoming and/or outgoing directed edges are re-directed to point from or to different nodes (Fig 3, top-right). Independent assortment during meiosis can be modeled via edge-mixing of parental GRNs such that an offspring acquires alleles, i.e., edges in the GRN, from both parents (Fig 3, bottom). Similar to mutation, recombination is an edge-rewiring process that is constrained to swapping binding sites or transcription factors at the same locus. Finally, gene duplication is equivalent to adding a parallel edge that represents the identical allelic content of a duplicated locus.

**Figure 3:**
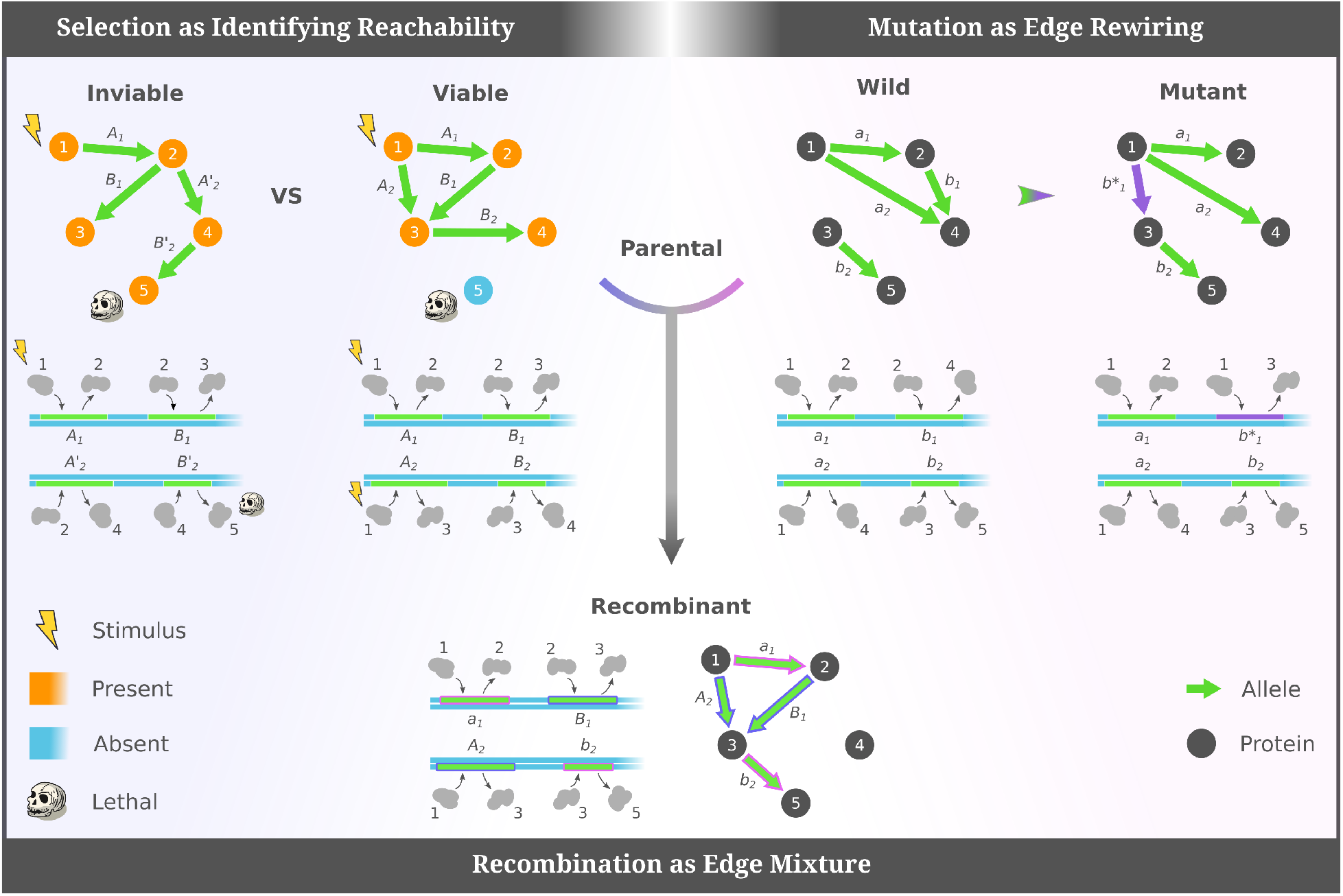
How the pathway framework turns evolutionary mechanisms into graphical operations on the GRNs. Since the pathway framework directly models the functionality of alleles of genes as edges, mutation, meiosis, and recombination can be modeled as edge-rewiring and edge-mixing, while a minimal selection scenario of binary fitness can be modeled as identifying “reachability” in a GRN.

An individual’s viability subjected to natural selection is a response to its molecular phenotypic status, which–under the pathway framework–can be modeled as a fitness function associated with the collective state of nodes and edges in the GRN. For example, one could study the time-varying concentration of each protein, attach a continuous dynamic or a stochastic reaction to every allele (Walczak et al., 2009), and/or define fitness as a function of the high-dimensional concentration vector, etc.. On the other extreme, we can consider Boolean networks, which have been shown to effectively capture many of the most relevant dynamical features of empirical regulatory systems (Davidich and Bornholdt, 2008). In this minimal scenario, each protein is assigned to a Boolean state (present or absent) and external environmental signals stimulate the existence of specific proteins in the organism. The logical states then cascade through the genetic pathways, where–given the presence of a gene’s transcription factor–loci activate and generates protein product(s). The phenotype of a GRN is thus the “reachability” from the environmental stimuli, whose binary survival is defined via a sharp fitness landscape over plausible collective Boolean states (Fig 3, top-left).

We adopt the Boolean-state assumption of GRNs because they readily shed light on the formation of hybrid incompatibilities. Hybrid incompatibilities are lethal combinations of alleles that were not prevalent or present in parental lineages, but are in hybrids. Moreover, the combination is minimal in the sense that the lack of any of its allelic elements will not lead to an inviable hybrid. In the pathway framework, suppose that binary viability only depends on a set of lethal proteins, i.e. an individual will not survive selection if any of those protein are present, a combination of alleles that includes a pathway from a environmental stimulus to a lethal protein makes the GRN inviable. If the alleles exactly comprise a simple path, which contains no cycles, they become a minimal combination and thus form an incompatibility. Additionally, The complexity of genetic interactions can be characterized by the number of alleles involved, which is called the order of hybrid incompatibility and related to the length of the simple pathway (in particular, *n* + 1 alleles form an incompatibility of order *n*).

### Simulating the evolution of GRNs

Briefly, we consider a Wright-Fisher model of evolution with natural selection, i.e., constant population size, no mutation, no migration, non-overlapping generations, and random mating. Selection occurs during the haploid stage of the life-cycle, where individuals that survive selection fuse randomly, i.e., create diploids, and undergo meiosis to generate the subsequent generation. Populations are seeded such that each individual has a randomly generated GRN and evolve until a single GRN fixes in the population. Simulations are further detailed in the Methods.

Fig 4a shows the proportion of individuals in the population that survive natural selection. Initially, due to the variation of randomly seeded GRNs, the fraction of viable individuals differed substantially between simulations with different initial conditions. However, as the gene networks evolved, the population’s viability increased and quickly reached a state where every individual survived selection (dashed line). During this 100% survival stage, natural selection was no longer effective and the population evolved to fixation via genetic drift. Not surprisingly, our results demonstrate that GRNs can rapidly evolve from a heterogeneous population with low average viability to “match” an imposed selective regime or environment.

**Figure 4:**
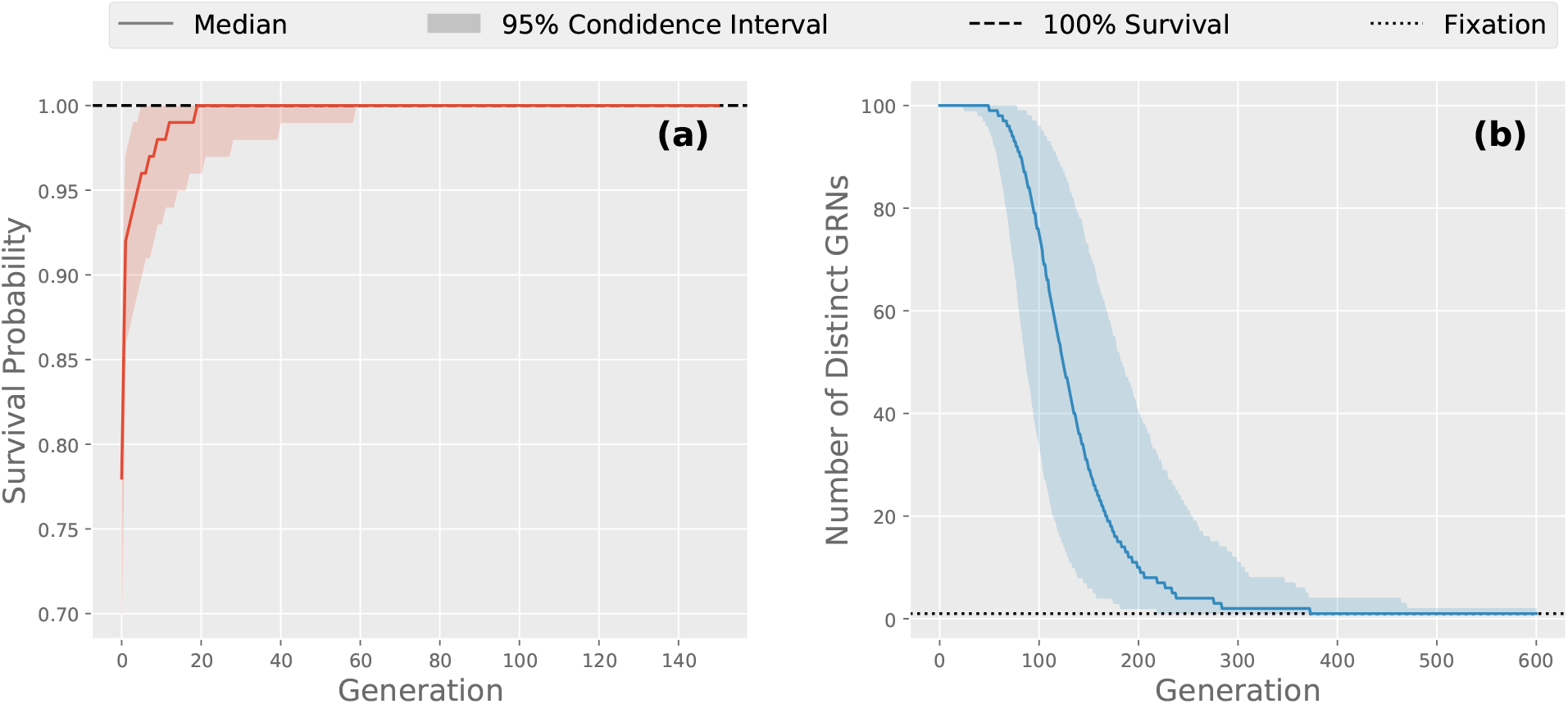
Populations adapt to the environment and then fix a single GRN. Here, we show for every generation of GRN evolution, across multiple allopatric populations with different initial conditions: **(a)** the survival probability of an individual and **(b)** the number distinct GRNs in each population, where two individuals’ GRNs were deemed effectively identical if they were isomorphic. The average viability of each population increased over time and rapidly achieved 100% survival, which indicates that evolution of GRNs drove adaptation toward the imposed environment. We also observe decreased variation of GRNs as they evolved, with individuals in the same allopatric population, i.e., simulation run, eventually fixing for the same GRN.

In addition to achieving 100% survival, populations always fixed a single GRN. Fig 4b plots the number of structurally-distinct GRNs in each generation. The decreasing trend demonstrates that, although various GRNs have equal survival probability, it becomes more and more likely that individuals shared a common GRN. Moreover, the populations always fixed a single GRN (dotted line) after a sufficiently long period of time. This phenomenon can be intuitively explained by the mechanism of sexual reproduction. In our model, parents with identical GRNs would lead to offspring of the same GRN, since any two corresponding groups of segregated alleles retrieved the parental gene network. Thus once there was a majority GRN in the population, it would have a higher chance of retaining its genetic configuration in the next generation, as compared to being replaced by meiotically shuffled variants.

Lastly, to better understand how parallel lineages evolve, we consider a scenario where multiple allopatric populations are seeded with the *same* initial conditions. Similarly, each allopatric population rapidly achieved 100% survival and then fixed a single GRN. However, across allopatric populations seeded from the same initial conditions, many different GRNs fixed. Fig 5 presents the distribution of fixed GRNs for a smaller-scale simulation (Setup 2 in Methods). We see that the fixed GRNs were diverse and non-uniformly distributed. Despite being under identical selection forces and having the same initial condition, lineages evolving from a common ancestral population fixed alternative GRNs. This result demonstrates that a broad range of GRNs can survive the given selection pressure. Furthermore, none of the viable GRN structures had a zero fixation probability, indicating a thorough exploration of evolution in the space of possible GRNs. That so many different GRNs fixed suggests that evolution was less governed by a definite trajectory, but instead it occurs via an uncertain realization among all the possibilities constrained by the ancestral population and the selection pressure.

**Figure 5:**
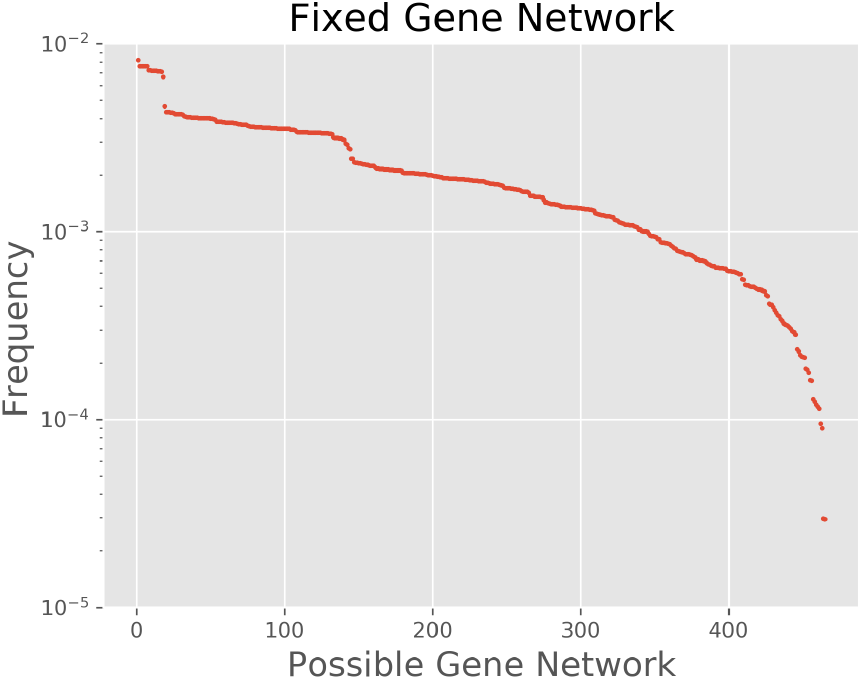
Fixation of parallel lineages resulted in a wide range of GRN structures. We simulated isolated populations from the same initial conditions until they reached fixation. In this case Setup 2 in Methods was applied in order to tractably enumerate all plausible GRN, and the ancestral populations were chosen such that the fixation was unbiased by the initial allele frequencies. The 10^7^ acquired GRNs were categorized into 465 viable structures and the fixation frequency of each structure was plotted in a descending order. The distribution shows that isolated lineages fixed alternatives gene networks, some among which were more favorable under our model of GRN evolution.

### Reproductive barriers arose rapidly as gene networks evolved

If the survival probability and fitness of GRNs were identical, the distribution of fixed networks should be uniform over all viable conformations. Because we observe a strongly non-uniform distribution (see Fig 5), some other form of selection, i.e., as opposed to simply viability selection, is likely operating on the GRNs. We note that during random mating, even between two parents with viable GRNs, some of their shuffled offspring can be inviable. Coupled with the observation that different allopatric populations, i.e., simulation runs, fix alternative GRNs from the same initial conditions, we hypothesized that some degree of reproductive isolation may exist between these fixed populations.

To test for the presence of reproductive isolation, we performed a “hybridization” experiment between parallel lineages that had reached fixation. Starting with lineages branched from a common ancestral population, two fixed lineages were randomly selected and interbred. Hybrids were generated and the reproductive isolation metric (RI) between the parental populations was computed (see Methods). By repeating this procedure, we obtained a distribution of reproductive isolation, as demonstrated in Fig 6a inset. Despite a large fraction of crosses resulting in nearly zero RI, we discovered pairs of lineages with positive reproductive isolation metric. Specifically, the RI distribution displays several regions of positive reproductive isolation such that a higher percentage of hybrid offspring are inviable. Positive RI was also observed in simulations with larger population size, although less frequently (see S1 Fig). Thus, we conclude that reproductive barriers between fixed lineages, derived from the same initial population and experiencing identical selection, occur even when the effects of genetic drift are substantially reduced.

**Figure 6:**
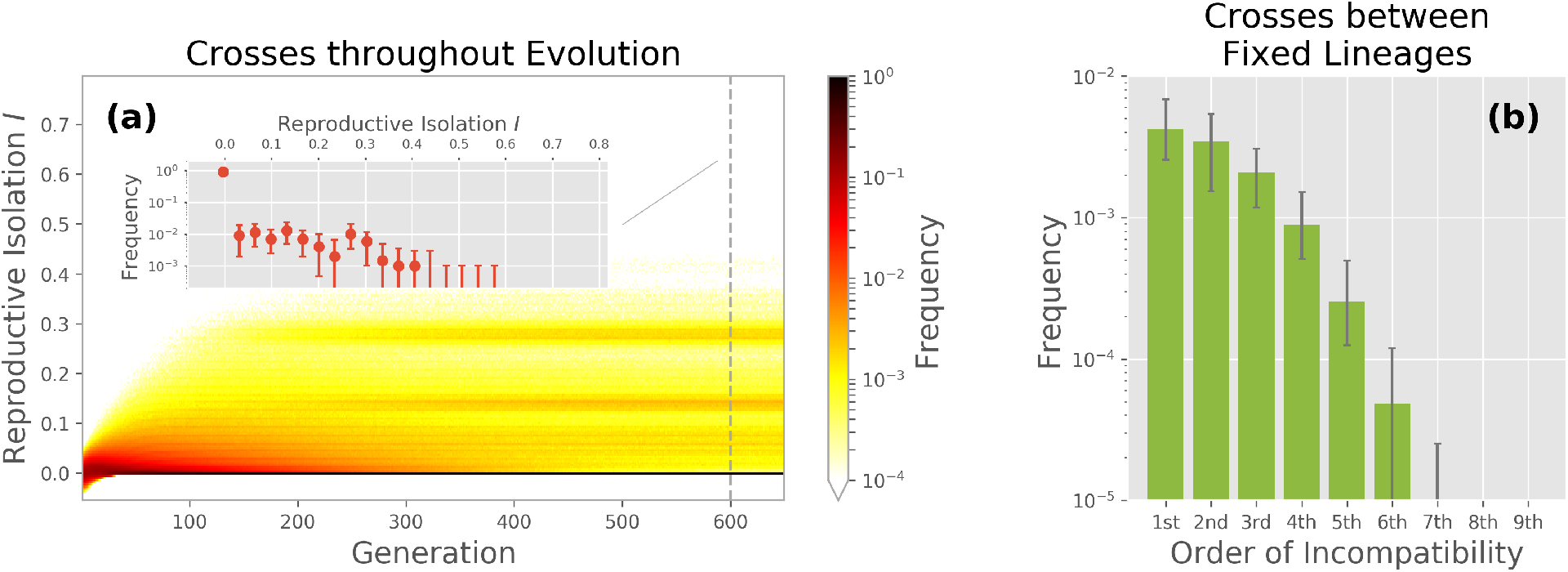
Reproductive barriers arose rapidly between allopatric populations. **(a, Inset)** Distribution of reproductive isolation between pairs of fixed lineages. A non-negligible fraction of crosses led to positive reproductive isolation, which reflects the occurrence of inviable hybrids and indicates reproductive barriers between fixed lineages. **(a)** We crossed allopatric populations at every generation during GRN evolution and stacked the RI distributions into a heat map. A vertical slice in this heat map represents the RI distribution at a given time, similar to the inset, but where the color shows the mean frequency for each bin. The growing level of positive RI indicates that reproductive barriers arose at the early stage of evolution. **(b)** Frequency that incompatibilities with various order were observed among hybrids between fixed lineages. We see that the order of incompatibilities included a broad range and that the simple pairwise interaction did not significantly dominate over more complex incompatibilities. Moreover, hybrid incompatibilities are consistent with the clustered level of RI and hence sheds light on the observed RI distribution (S1 Appendix). In both the inset and panel (b), the plots show the statistic of the distribution among multiple groups of allopatric populations, specifically the median frequency and the 95% confidence interval.

Given noticeable reproductive barriers between fixed lineages, we further studied when those barriers first manifested during GRN evolution. Note that because our simulations did not contain mutation, incompatibilities arise because of shuffling during meiosis. Here, instead of waiting until GRN fixation, we instead evolve lineages for *T* generations and then cross them to generate hybrids as described above. By varying *T*, a series of reproductive isolation distributions were acquired. Fig 6a collects and displays them in a heat map. A vertical slice represents a RI distribution as in the inset panel, but crosses were made after *T* generations rather than waiting for lineages to reach fixation. We see that the regions of high incompatibility noted in Fig 6a inset becomes bands in the heat map, which allows us to trace the emergence of reproductive barriers.

Initially the reproductive isolation distribution was relatively symmetric around zero. However, As GRNs evolved, the range of RI broadened and its extreme value in the positive tail increased. The trend towards higher levels of RI decelerated after 100 generations; it then stabilized and formed a band structure, where crosses cluster around certain levels of reproductive isolation. Fig 6a hence reflects that reproductive barriers existed at low levels as soon as the lineages started evolving independently and peaked at a time prior to GRN fixation. Because, by assumption, the model does not include mutation, all of the alleles underlying RI were present in the ancestral population. Nevertheless, we observe that RI between isolated populations peaked well before fixation of GRNs.

Next, for incompatible hybrids generated in our crossing experiment, we determine how complex the underlying mechanism of RI was. Specifically, Fig 6b shows how frequently an inviable hybrid resulted from an incompatibility of a certain order. We see that hybrid incompatibilities spanned a broad range of interaction orders. Importantly, the simple two-allele interaction was only slightly more common than incompatibilities resulting from three or four interacting alleles and interactions above forth order made up almost 3% percent of all incompatibilities. However, we note that the frequencies of incompatibility order varied depending on the ancestral population.

The pattern of complex genetic interactions provides insight into the distribution of reproductive isolation. Based on the independent assortment mechanism in our model–and assuming that multiple incompatibilities rarely occurred between two parental GRNs–we conclude that hybrid incompatibilities quite often involved higher order interactions, which did not arise as a result of selection, but simply were an expected consequence of GRNs being high order (S1 Appendix). Further, the discrete characteristic of hybrid incompatibilities led to a higher likelihood at certain RI levels. The band structure in Fig 6a agrees with this prediction (S1 Appendix), which suggests that reproductive barriers are strongly influenced by the concealed hybrid incompatibilities and are coupled with the genetic interaction pattern shown in Fig 6b.

### Early divergence between lineages was critical for reproductive barriers to emerge

To further study the emergence of reproductive barriers in our model, we investigated the relative importance of various evolutionary forces in generating the observed patterns of RI. In particular, were the barriers attributed to selection pressure, random genetic drift, or both? We designed two “control scenarios” that were based upon the previously simulated model, but contained modifications to remove the effects of either selection or drift. Comparing the strength and pattern of RI resulting from the two control scenarios, i.e., the removal of drift or selection, to the original GRN dynamics, which contain both evolutionary forces, provides an assessment of the removed component’s role in shaping the observed pattern of RI.

Removing the effect of natural selection is straightforward to simulate. In this control scenario, populations simply evolve in a selectively neutral environment where all GRNs are viable. Thus, all individuals survived and genetic drift became the only remaining evolutionary force. Of course, this neutrality concurrently made the RI metric ill-defined. We avoided this issue in the crossing experiments to calculate RI by placing the parental populations under the same non-neutral environment in the original model, so the hybrids would be generated from survivors subjected to selection pressure. The reproductive isolation metric could then be computed with respect to the non-neutral environment. Placing the parental population through a round of viability selection just prior to hybridization ensures comparability between the model and the “no selection” control scenario since the survivability of hybrids was evaluated under the same environment and was not biased by the otherwise inviable parents.

Fig 7a shows the contrast of barriers observed in the original GRN evolution model (red) and in the scenario with no selection (blue). We traced a measure of reproductive isolation over time, defined as the 99th percentile of the RI distribution, which is a sufficient indicator of reproductive barriers between lineages. We discovered that in both the model and the control scenario, the leading RI *I*\* increased and then saturated. Furthermore, the growth in *I*\* decelerated after a similar number of generations in both scenarios. That RI occurs at a higher level in the control experiment indicates that selection did not “cause” the fixation of barriers between allopatric populations, but instead suggests that selection was actually limiting chances for incompatibilities to occur in hybrids. We hypothesize that–although restricted as compared to drift–selection operating on incompatibilities likely induced the observed disconnect between viability and fitness seen in Fig 5.

**Figure 7:**
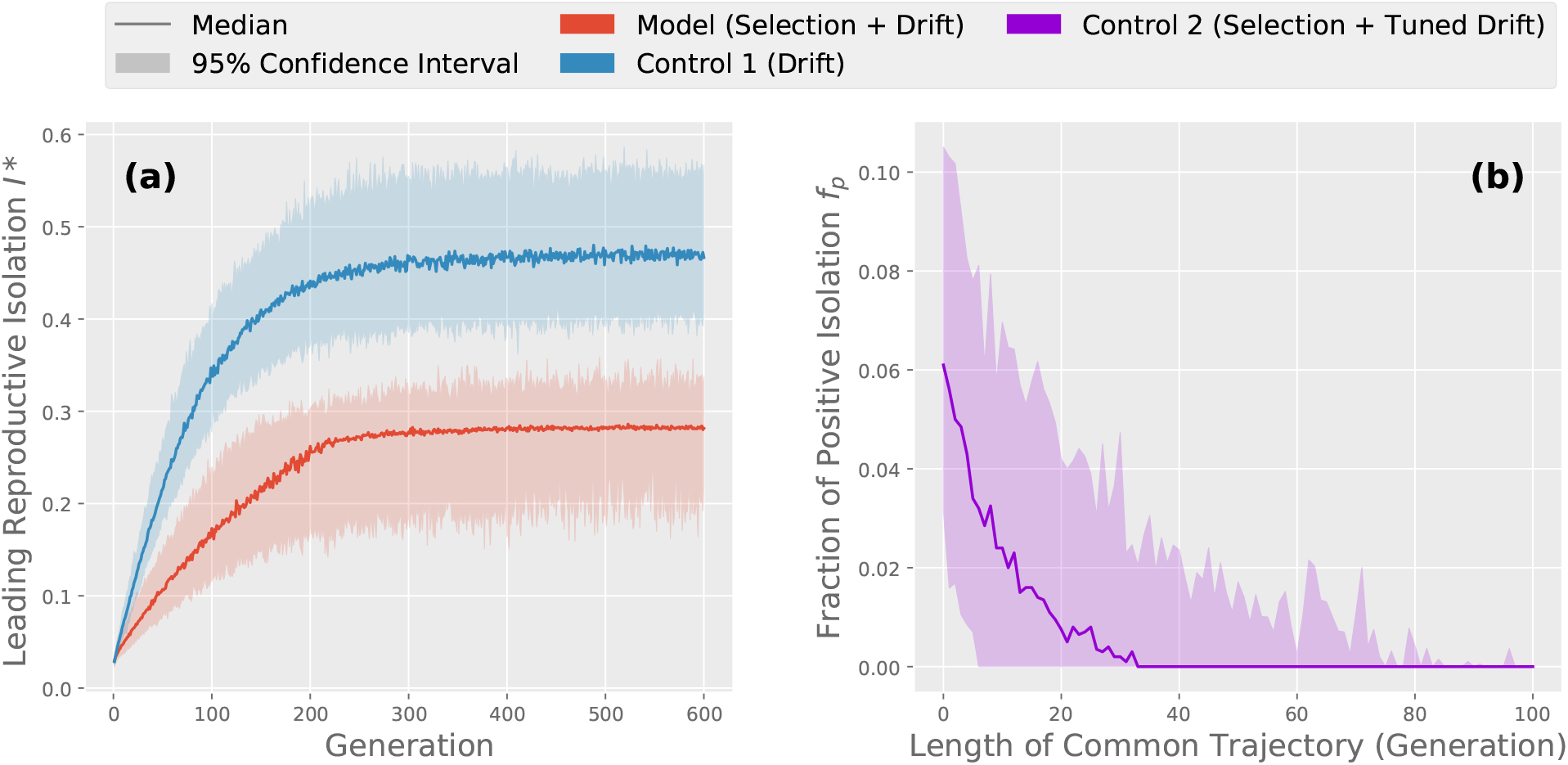
Early divergence of evolutionary trajectories between lineages was necessary for reproductive barriers to arise. Here we compare a statistic, termed leading reproductive isolation *I*\* (99th percentile of the RI distribution), measuring the degree of reproductive barrier in the original model and two designed control scenarios. Control scenarios were simulated with the same group of ancestral populations as the model, where lineages were then crossed to generate hybrids. **(a)** Leading reproductive isolation *I*\* among allopatric populations over time, where positive values indicate the existence of reproductive barriers. We plot the original model in red and the control scenario with a neutral environment in blue. The increasing and larger *I*\* uncovered in the control scenario implies that reproductive barriers were still observed when the selection forces were silenced. **(b)** Long-term fraction of positive RI *f_p_* when the influence of random genetic drift was tuned. We simulated the evolution of lineages, but first confine them to a common trajectory of length *L*, which was realized by evolving a single population from the ancestors for *L* generations, and then simulated allopatric evolution from this now less diverse ancestral population. The original model corresponds to the case where *L* = 0, and for any positive *L* the effect of drift were lessened. We obtained the *f_p_* metric when lineages evolved for 600 generations, where *f_p_* = 0 suggests no barriers among populations. That *f_p_* decreased with *L* to 0 shows that reducing the effect of drift diminished reproductive barriers. As a result, it implies the criticality of divergence among evolutionary trajectories for barriers to emerge.

We next turned to the contribution of genetic drift. This control scenario, however, was less straightforward due to technical difficulties associated with directly removing random genetic drift from the model. Neither abandoning sexual reproduction nor simulating an infinite population would result in non-trivial and/or computationally tractable GRN evolution. Alternatively, we designed a control scenario where the evolutionary influence of drift could be tuned and limited. Genetic drift results in stochasticity and causes populations to experience diverse trajectories. On the other side of the coin, if two lineages show similar evolutionary trajectories, one would say that drift effectively leads to less divergence between them. We restricted the influence of genetic drift by first confining lineages in a common trajectory for *L* generations, and then freed the populations and let them evolve independently, i.e., in allopatry. Varying the length of the common trajectory *L* tunes the overall similarity among lineages. Therefore, *L* quantitatively reflects the strength of genetic drift.

Fig 7b demonstrates the long-term fraction of positive reproductive isolation introduced in Methods, termed *f_p_*, as we varied the length of the common trajectory. Despite substantial variation in *f_p_* in the original model, which corresponds to the case where *L* = 0, a decline of *f_p_* was uncovered as early evolutionary confinement was extended. We discovered 50% of the experiments showed a zero *f_p_* after lineages were evolved together for 40 generations, and as the length of common trajectory exceeded 80 generations positive reproductive isolation was hardly found between lineages. More importantly, Fig 7b suggests that as the evolutionary influence of genetic drift was mitigated, RI was weakened and eventually vanished. Namely, restricting early divergence among populations due to genetic drift diminished reproductive barriers. This control scenario consequently suggests that divergence between lineages, coupled with high diversity in the ancestral population, is critical for reproductive barriers to arise. That RI is still present, despite substantial shared evolutionary history, supports our earlier finding that positive RI was also observed in simulations with population sizes that were an order-of-magnitude larger (see S1 Fig).

### Intra-lineage incompatibilities were eliminated stochastically while inter-lineage incompatibilities persisted and led to reproductive barriers

To better understand how reproductive barriers might be removed within a lineage, but persist between lineages, we computed two quantities from the underlying genetic pool. First, the size of the genetic pool, which determines how many possible genotypes a population contains. This measure captures the potential genetic diversity in the population. Second, we count the number potential incompatibilities in the underlying genetic pool, which are lethal allelic combinations that could potentially be realized in the next generation. These incompatibilities compose the source of inviable offspring and RI between allopatric populations. However, because even for small GRNs searching for all possible incompatibilities quickly becomes computationally intractable, we developed a novel algorithm (summarized in Methods) to compute their number in the genetic pool.

Because our model does not contain mutation, one would expect the size of the underlying genetic pool to decline in our simulated gene network evolution. Any allele in an individual was inherited from its parents, and thus it must appear in the parental generation as well. Additionally, a parental allele might not persist in the offspring for two possibilities: either it was not transmitted because of finite population size of the progeny generation and the stochasticity during sexual reproduction, i.e. drift, or it formed a lethal pathway along with other inherited alleles which made the offspring inviable, i.e. selection.

Fig 8a demonstrates the size of the genetic pool over time, where we compare simulations in the original model (red) and in the control scenario without selection pressure, i.e., only genetic drift will reduce the size of the genetic pool (blue). A rapid decline of genotypic diversity was witnessed under both models. More intriguingly, little difference was found between the GRN evolution model and the control scenario under a neutral environment. The two median curves nearly overlap, and for any given generation, the pool size in the original model was not significantly smaller than the control counterpart. Therefore, we find additional support for our earlier finding that–although both natural selection and random genetic drift decreased genotypic diversity–drift was the dominant evoluiontary force. Nevertheless, while the effect of drift reduced diversity within a lineage, it (again as expected) increased the divergence between lineages.

**Figure 8:**
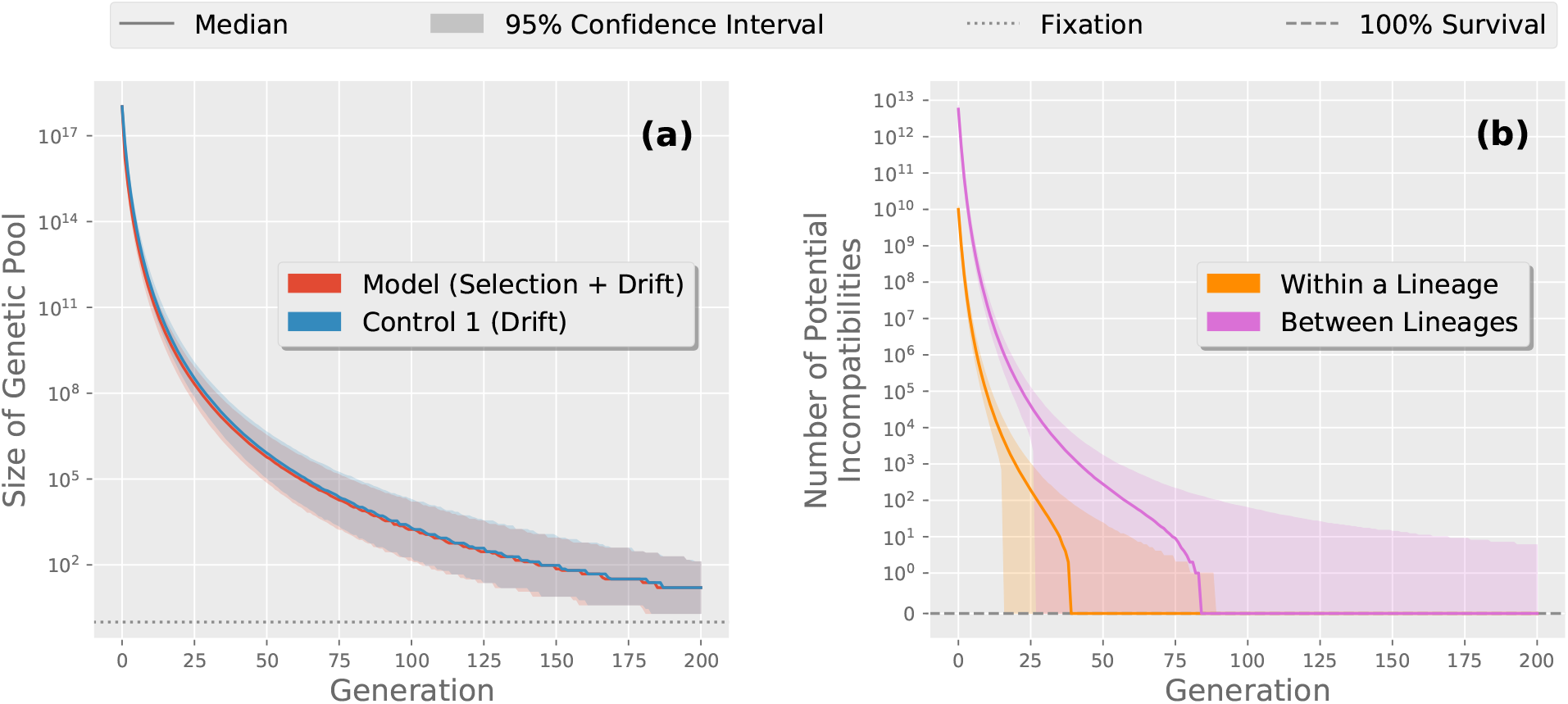
The underlying genetic pool lost alleles and eliminated potential incompatibilities within allopatric populations, whereas inter-lineage incompatibilities persisted. **(a)** Size of the underlying genetic pool for each generation, where we plot the original model in red along with the no selection control scenario in blue. Both cases show a similar reduction in the genetic pool. The similarity of these curves suggests that the continual losses of allelic diversity within a lineage was dominated by random genetic drift. **(b)** Number of potential intra-lineage (orange) and inter-lineage (pink) incompatibilities for each generation in the original model. We found that the number of potential incompatibilities also decreased as GRNs evolved, which is explained by the reduced allelic diversity in the genetic background. The vanishing intra-lineage incompatibilities implies disappearing sources of inviable hybrids, and it provides a mechanistic understanding of how a genopytically rich populations adapted to the imposed environment. Contrarily, the intra-lineage incompatibilities remained during GRN evolution. It was the persistent potential incompatibilities between allopatric populations that led to evident reproductive barriers.

Fig 8b shows the number of potential incompatibilities within a lineage’s underlying genetic pool (orange). We found that the number of incompatibilities embedded in a population also decreased over time. This phenomenon is understood by the continual loss of allelic diversity, since removing an allele from the underlying pool always restricts the possibilities to form a lethal pathway in the GRN. Furthermore, the number of potential incompatibilities fell rapidly until no potential incompatibilities remained. The elimination of potential incompatibilities illuminates how a population adapted to the imposed environment when GRNs evolved, as shown in Fig 4a. Random genetic drift drove the loss of a lineage’s genotypic diversity, and along with the guidance of selection, it eliminated probable lethal pathways in the genetic background. Once all the potential incompatibilities were eliminated, no source of inviable offspring existed and consequently the population reached 100% survival. Again, this result supports our earlier finding that natural selection was operating against incompatibilities within a lineage, but that drift was nevertheless the dominate force in structuring incompatibilities between lineages.

Finally, we investigated incompatibilities between underlying pools of lineages, which we call the “inter-lineage” incompatibilities, as compared to potential lethal allelic combinations within a population termed “intra-lineage” incompatibilities. Fig 8b presents the number of inter-lineage incompatibilities over generations (pink). We observed more incompatibilities between allopatric populations than those within a population, and similarly their amount dropped as allelic diversity decreased. In contrast, inter-lineage incompatibilities were removed at a slower pace compared to intra-lineage incompatibilities. The sustained confidence interval further suggests that some inter-lineage incompatibilities persisted, which was also the case after populations reached fixation (S2 Fig). The persistence of these potential incompatibilities qualitatively explain the inviable hybrids revealed after GRN evolution. In spite of lineages adapting to the same imposed environment, hybrdiziation can “resurrect” a lethal combination of alleles, which was eliminated in either lineages yet remained in their joint genetic background. This explanation also supports the stronger barriers uncovered in the neutrally evolving control in Fig 7a, since inter-lineage incompatibilities would be more persistent without the constant selection pressure (S2 Fig).

## Discussion

In this work, we develop a pathway-oriented construction of GRNs where alleles are represented as edges in a network. Termed the pathway framework, this model allows us to apply network science analyses to the study of speciation. Specifically, we simulate the evolutionary dynamics of GRNs under a model that includes natural selection, sexual reproduction, and genetic drift. Starting from a diverse ancestral population, we show how reproductive isolation can arise rapidly between allopatric populations experiencing identical selection pressure, which agrees with empirical studies recently reviewed in Coughlan and Matute (2020). Then, using a series of counter-factual simulations, we disentangle the relative importance of each evolutionary force included in our model and identify the central roles of high-dimensionality and functional redundancy, even in comparatively small GRNs, for speciation. Finally, we show how higher-order genetic incompatibilities can often evolve simply as a by-product of GRN evolution.

Our counter-factual simulations reveal that the observed reproductive barriers likely resulted from divergent evolutionary trajectories and persistent, inter-lineage incompatibilities. Driven by genetic drift and guided by selection, many GRNs that satisfied the same viability function were sorted into parallel lineages, whereas mixing edges between them can lead to fatal pathways and inviable offspring. These results highlight the importance of “functional redundancy” in evolution (Nowak et al., 1997; Láruson et al., 2020) and agree with earlier studies that suggested alternative regulatory structures can achieve the same phenotype (True and Haag, 2001; Wagner and Wright, 2007; Schiffman and Ralph, 2018; Tkačik and Walczak, 2011). Indeed, both theoretical and empirical studies increasingly support the role of parallel trajectories through fitness landscapes in evolution (Elmer and Meyer, 2011; Bank et al., 2016; Ogbunugafor and Eppstein, 2016; Langerhans, 2018).

More importantly, the pathway framework illustrates why degenerate genotypes can reach fix through parallel evolution. Once the alleles are presented as functional pathways connecting an underlying group of proteins, the conjunction between genetic factors and physiological traits is no longer a bipartite mapping; the phenotype, as the collective chemical status of proteins, is a convolution of active signals and external stimuli propagating on the network of genetic pathways. The pathway configuration that satisfies a specific environmental input and phenotypic output is, as a result, not unique. One can thus find numerous, functionally degenerate gene network structures fulfilling the input-output viability relation, as Fig 5 demonstrates. In addition, taking advantages of basic network analyses, the pathway framework predicts that the number of GRNs generating the same phenotype will increase more than exponentially as the system scales (S2 Appendix).

The minimal model of GRN evolution we consider encapsulates selection through binary viability, which is essentially a special case of holey adaptive landscapes (Gavrilets, 1997). Gavrilets and Gravner (1997) introduced a multi-locus model where each genotype was independently assigned to one of two fitness levels, whose results suggested that reproductive isolation can arise simply due to the high dimensionality of the genotype space. In a similar vein, our model further connects the high dimensionality of genotypes to complex genetic interactions. Under the pathway framework, inviability originates via the mechanism of hybrid incompatibilities, i.e., allelic combinations that form lethal pathways in a GRN. Furthermore, the pathway framework can be readily extended to include alternative fitness landscapes. For example, Barton (2001) demonstrated that stabilizing selection can generate reproductive isolation, and the pathway framework can be easily embedded into such a continuous fitness landscape.

Our work supports the latent connection between speciation processes and ancestral genetic variation. Ancient polymorphisms drive genomic divergence and confound inference of evolutionary processes (Guerrero and Hahn, 2017). Additionally, these same polymorphisms–and the empirical evidence that incompatible alleles often far predate speciation events–have recently been consolidated into a “combinatorial” view of speciation (Marques et al., 2019). The combinatorial mechanism proposes that, if there was a past admixture event or if standing genetic variation persists, the reassembly of these old genetic variants can facilitate rapid speciation and adaptive radiation. Marques et al. found that ancestral genetic variants that had undergone selection–and thus are likely to be beneficial–often have higher allele frequency than *de-novo* mutations. Alternatively, we demonstrate that stochastic loss of accessible pathways resulted in the fixation of incompatible GRNs due to their functional redundancy and high dimensionality. We also observed that the emerging reproductive barriers required the ancestral variation to be greater than a critical amount (S3 Appendix). Our pathway framework hence adds theoretical support for the role of stable polymorphisms in hybrid incompatibilities, as reviewed in Cutter (2012); Maheshwari and Barbash (2011). We therefore consider the evolution of regulatory pathways as a parallel mechanism with which ancestral genetic variation can facilitate the appearance of new species.

Recent evidence supports our findings that distributed regulatory networks are sources of genetic incompatibilities between closely related taxa. For example, Morgan et al. (2020) identified a number of disrupted gene expression modules in sub-fertile, hybrid mice and concluded that “hub” genes in these modules played a central role in genetic incompatibility. Additionally, Rougeux et al. (2019) showed how gene expression was disrupted in hybrids between benthic and limnetic species pairs of Lake Whitefish (*Coregonus clupeaformis*) and that genes underlying this disruption were enriched for polymorphisms in the outgroup taxa, the European Whitefish, *Coregonus lavaretus*. Similar patterns have also been found in repeated adaptation to freshwater, benthic envionrments in the threespine stickleback (*Gasterosteus aculeatus*) (Erickson et al., 2016). This pattern of gene network disruption and standing genetic variation is consistent with our findings from the pathway model. Furthermore, Guerrero et al. (2016, 2017) found evidence for the role of gene regulatory disruption and the presence of persistent antagonistic interactions in speciation in *Solanum*. Lastly, Stankowski et al. (2019) found that genetic divergence arose rapidly after population of monkeyflowers were isolated and that the evolution of regulatory-based genetic incompatibilities may have been driven parallel selection pressure from a polymorphic ancestor (Stankowski et al., 2019; Jiggins, 2019), again the mechanism identified in the pathway framework.

Our work is not without important caveats and there are many clear opportunities to advance the pathway framework. First, our model did not include mutation, large-scale genome rearrangements, nor whole genome duplication events, which are all known to be important for genetic incompatibles and speciation (Otto and Whitton, 2000; Noor et al., 2001; Kirkpatrick and Barton, 2006; Hoffmann and Rieseberg, 2008; Guerrero et al., 2012). Although it is possible to draw some preliminary conclusions regarding the effect of random mutations from our counter-factual simulation that “eliminated” genetic drift, we leave a fuller exploration of mutation for future work. Second, despite the widely documented, asymmetric risk of hybrid breakdown in the heterogametic sex, i.e., Haldane’s rule (Haldane, 1922; Coyne and Orr, 1997; Delph and Demuth, 2016), our model considers sexual selection with only a single sex of mating type. Third, both empirical results from yeast (Bernardes et al., 2017) and from theoretical, population-genetic models (Dagilis et al., 2019) point towards the importance of increased hybrid fitness, i.e., heterosis, even if only temporary, during speciation (Gavrilets, 2003). Forth, there are studies that clearly demonstrate the importance of divergent selection in the process of speciation (e.g., Nosil et al., 2002; Allender et al., 2003; Gow et al., 2007). However, the pathway framework can be readily modified to include divergent selection and will almost certainly result in higher degrees of reproductive isolation. Fifth, recent work suggests that demographic complexity such as a varying population size can adds constraints on the process of speciation (Harvey et al., 2019; Khatri and Goldstein, 2019; Yamasaki et al., 2020; Momigliano et al., 2020). Future research may combine the pathway framework with additional demographic information to further investigate how ancestral genetic variation is sorted throughout the evolutionary trajectories. Finally, the relative importance of post-zygotic, genetic incompatibilities in generating and/or maintaining species remains an active area of investigation and scientific debate (Servedio and Sætre, 2003; Rundle and Nosil, 2005; Rieseberg and Willis, 2007; Magnuson-Ford and Otto, 2012; Hopkins, 2013; Seehausen et al., 2014). Our results support a growing body of literature on the theoretical importance of higher-order, genetic interactions in the speciation process (Johnson and Porter, 2000; Palmer and Feldman, 2009; Schiffman and Ralph, 2018; Blanckaert et al., 2020) and are consistent with emerging empirical data on genes involved in reproductive isolation (Seehausen et al., 2014; Marques et al., 2019; Coughlan and Matute, 2020). We support calls for the increased use of high-fidelity simulation models in evolutionary genetics (Jiggins, 2019; Satokangas et al., 2020), but stress the need for models with interpretable mechanisms and that generate testable hypotheses (Shou et al., 2015). For example, our results on the evolution of higher-order incompatibilities could serve as a null model for evaluating empirical data under the relaxed assumption that genes function independently. Only by joining mathematical and computational theory with comparative-level data can we uncover general patterns in speciation and, potentially, resolve long-standing debates in the field.

## Methods

### Numerical simulations

#### General schema and assumptions

In this work we simulated evolution GRNs in allopatric populations. Throughout evolution, we assumed that individuals had a constant number of loci and thus a fixed number of edges in their GRNs. The underlying set of nodes in GRNs also remained unchanged as we reasoned in Results. We further introduced different categories of nodes/proteins to concrete the space of plausible alleles. Some proteins were presumed to only be present with the environmental stimuli, which were not products of any locus; on the other hand, some other proteins were presumed to have mere physiological effects, and thus they were not capable of activating gene expression. We called them source proteins and target proteins respectively. A plausible allele was therefore labeled by a non-target protein that could activate its expression and a non-source protein that would be synthesized. In our simulations we supposed only one source protein and one target protein.

We considered a naive model of GRN evolution incorporating natural selection, independent assortment and random genetic drift. The environmental condition was set fixed over time and across populations. We assumed that the environment stimulated presence of one protein and it specified another protein with a lethal effect (specifically, they reconciled with the source and the target protein respectively). Viability of individuals was presumably equated to the reciprocal binary state of the lethal protein. Hence given the current generation, individuals were selected such that whoever did not possess a pathway from the environmental stimulus to the lethal protein survived and were able to reproduce.

The survivors then randomly mated and formed the next generation with independent assortment. Here we assumed individuals with haploid-dominant life cycles, where the multicellular haploid stage is evident. During reproduction, specialized haploid cells from two individuals combined and formed a diploid zygote. The zygote experienced meiosis and generated haploid spores, which then developed into multicellular-haploid-stage individuals through mitosis. Supposed even segregation during meiosis of the diploid zygotes, we modeled the process of independent assortment as follow. Two parental individuals were randomly sampled from the survivors. The set of loci was first randomly partitioned into two groups of equal sizes. The offspring inherited alleles of one group of loci from one of its parents and alleles of the remaining loci from the other parent. Hence half of the edges in the offspring’s GRN came from one parent’s GRN and the rest was acquired from the other. This procedure was repeated until the next generation had the same constant population size as their predecessors.

#### Simulations and parameter setups

Here we summarize the two different parameter setups in our simulations:

**Setup 1:** We assumed 11 possibly existing proteins in the organism. A generation was composed of 100 individuals with 10 loci each. We generated 100 ancestral populations where individuals’ GRNs were randomly sampled from all plausible genotypes. For every ancestral population, we in parallel ran 100 simulations from it, which were regarded as lineages evolving in isolated geo-locations.
**Setup 2:** We assumed 5 possibly existing proteins in the organism. A generation was composed of 16 individuals with 4 loci each. We generated 10^4^ ancestral populations induced from a genetic pool containing all plausible alleles for each locus (which are samples drawn from all possible populations that own the same underlying genetic pool). For every ancestral population, we in parallel simulated 10^3^ lineages from it.

The randomly generated ancestral populations encapsulate our assumption of ancestral genetic variation, which reflect divergence of gene regulation that has been found in empirical studies (Gould et al., 2018). Setup 2 aimed to examine how broadly, in terms of fixed GRNs, evolution can explore in all possibilities. Thus it consisted of a larger amount of simulations starting with unbiased ancestral populations that were induced from a maximal genetic pool. If not otherwise specified, simulations shown in Results were run under Setup 1.

When we inspected reproductive barriers between allopatric populations by interbreeding them, we first sampled 1000 pairs of lineages and then each generated *F*_1_ 1000 hybrids. The survival probability of hybrids can then be obtained for all crosses. The same sampling procedure was also applied when we computed the number inter-lineage potential incompatibilities between pairs of allopatric populations.

### Metrics of reproductive isolation

We introduce a quantitative measure of reproductive isolation between lineages which evolved from a common ancestral population. Given a group of lineages and a chosen pair among them, the reproductive isolation between the pair is defined as the relative difference of hybrid survival

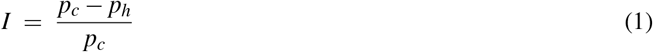

where *p_h_* is the survival probability of *F*_1_ hybrids, and *p_c_* denotes the average of survival probabilities of all lineages’ next generation. A positive value of reproductive isolation *I* implies that the hybrids have less survivability than the expectation of the offspring. In the extreme case where no hybrid lives, *I* = 1. It therefore serves as an indicator of reproductive barriers between two lineages.

Strengths of reproductive barriers among the group of lineages are described through a distribution of reproductive isolation, which can be obtained by sampling pairs of lineages and computing their reproductive isolation *I*. We further introduce two indicators for the existence of reproductive barriers. A quantity named leading reproductive isolation *I*\* is defined as the 99th percentile of the reproductive isolation distribution. It signals that there is one percent of crosses with reproductive isolation equal or larger than *I*\*. We would also like to raise a caveat that *I*\* > 0 is sufficient for the existence of reproduction barriers but not a necessary condition, due to the possibility of positive *I* in the distribution even if *I*\* ≤ 0. The leading reproductive isolation metric hence summarizes a high level of reproductive barriers that can be found among the lineages. On the other hand, the fraction of positivity in the reproductive isolation distribution serves as a necessity indicator for reproductive barriers, which we denote as *f_p_*. The zero-value of *f_p_* implies that none of the crosses generate inviable hybrids more than the anticipation of the offspring and thus the absence of reproductive barriers. Contrarily, a positive *f_p_* does not satisfy existence of barriers considering small reproductive isolation subject to noise. These two indicators are beneficial for us to identify the responsible part of the model to the observed evolutionary consequences.

### Potential incompatibilities within and between genetic pools

An intra-lineage incompatibility is a group of alleles in its genetic pool, each of a unique locus, that generates a lethal pathway. In our model those incompatibilities are the only source of inviability, and hence the number of potential incompatibilities provides information about reproductive barriers. Nevertheless, counting the number of potential incompatibilities within a genetic pool through a brute-force manner is computationally intractable. Here we suggest a relatively efficient algorithm when the total number of loci is small. Our strategy is to turn the task into solving a graph problem. The genetic pool can be transformed to an edge-colored network where nodes once more represent possibly existing proteins in the organism. The edges correspond to available alleles in the pool, which are colored by their according loci. A potential incompatibility then becomes a simple path from an environmental input signal to a lethal protein node, with an additional constrain that no edges on the path have the same color. We call such a path an edge-colorful simple path (ECSP).

The proposed algorithm, as demonstrated in Algorithm 1 in S4 Appendix, counts the number of ECSPs from the source nodes to the targets nodes by having agents propagate on the edge-colored network iteratively. An agents is capable of keeping information of the trajectory, including its current position on the network, the colors of edges it has traversed and the nodes that it has visited. In Algorithm 1, the NEW-AGENT procedure creates an agent instance given its position, visited colors and nodes accordingly. This trajectory information is also accessible fields of the agent instance. Initially we deploy one agent on each source node. At every iteration, each agent is substituted by all of its possible successors who are a hop away, such that the hop along with the agent’s memory obeys an edge-colorful simple path. Those successors can be deduced from the agent’s trajectory information as shown in Algorithm 2 in S4 Appendix. The cautiously-designed rule of agent propagation guarantees that the total number of agents locating on the target nodes at the *n*-th iteration equals to the number of the desired ECSPs of length *n*. Moreover, since the order of an potential incompatibility is bounded above by the number of genes in the organism, iterations as many as the amount of edge colors in the network are sufficient to obtain a computationally feasible count of all potential incompatibilities. The efficiency of the algorithm can be further improved by, instead of keeping track of numerous agents, monitoring the distribution of agent states over iterations.

The same algorithm can be applied to count the number of inter-lineage incompatibilities as well. In this case the underlying genetic pools of both lineages are transformed into a single edge-colored network, whose edges then consist of alleles in the two pools and are again colored by their according loci. A ECSP on this composite network either only traverses through edges from one of the genetic pools, or it contains alleles from the two different pools. These two scenarios correspond to a incompatibility within and between genetic pools respectively. Therefore, by counting the number of ECSPs on the composite network, and subtracting by the number of potential incompatibilities within the two genetic pools separately, we can compute the number of incompatibilities between the two underlying genetic pools.

## Acknowledgements

Hold.

## Supplementary figures

**Fig S1:**
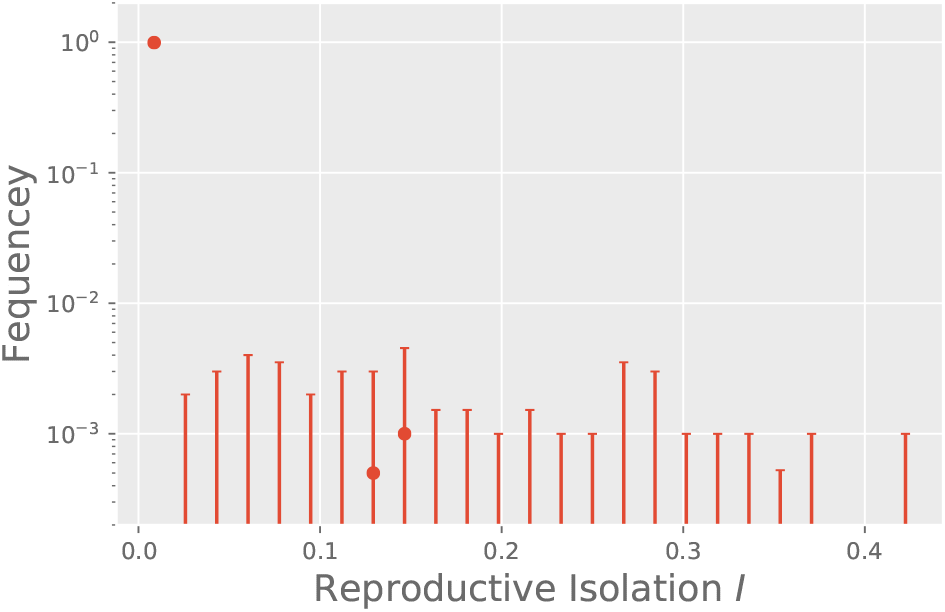
Weak RI appeared between fixed lineages of a larger population size. We also simulated the evolution of GRNs in allopatry with a larger population size, where a population contained 1000 individuals instead of 100 as in Setup 1 while other parameters remained the same. Here we show the distribution of RI between pairs of fixed lineages after the evolution for 6000 generations. Despite being less prevalent, inviable hybrids were still observed from the crosses, and the degree of RI was found to span through a wide range.

**Fig S2:**
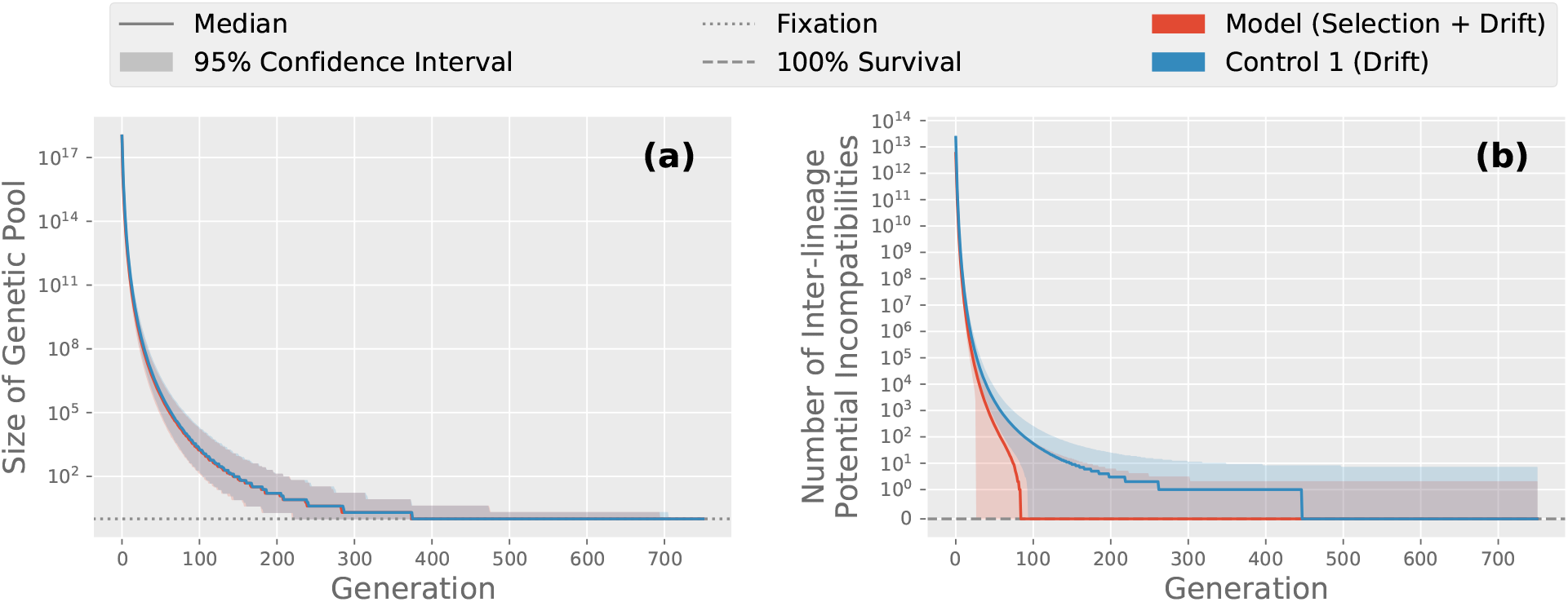
Inter-lineage incompatibilities were sustained throughout GRN evolution. **(a)** The size the underlying genetic pool continually shrank until there was only one accessible genotype. At this stage a population fixated a single GRN, and no significant difference was found between the model and the control scenario without selection, i.e., drift only. **(b)** In our model, inter-lineage incompatibilities persisted throughout evolution (red), which accounts for the sustained confidence interval of their abundance even after populations reach fixation. Interestingly, in the control scenario where natural selection was silenced, inter-lineage incompatibilities were eliminated at a slower pace. We hypothesize that due to the lack of guidance by selection, inter-lineage incompatibilities only became inaccessible through random genetic drift. This scenario led to fatal allelic combinations that were more persistent than those in the model and hence stronger reproductive barriers were observed.

## Appendices

## S1 Hybrid inviability against a single incompatibility

Here we analytically evaluate the probability that a hybrid is inviable presuming that multiple incompatibilities are rarely embedded in two parental gene regulatory networks. In addition, this naive analysis explains the pattern of RI distribution, Fig 6a in the main text.

Assume that there is only on incompatibility *ℐ* between the two parental gene networks *G*_1_ and *G*_2_. For convenience we suppose there are an even number of loci in the organisms, denoted by 2*m*, and let the incompatibility *ℐ* be of order *k −* 1 so it consists of *k* alleles to form a lethal combination. We also suppose that, among the *k* alleles in *ℐ*, *k*_1_ of them come from *G*_1_ and the other *k*_2_ alleles are from *G*_2_.

Following the rule of recombination between haploid GRNs in our model, the hybrid is generated by randomly segregating alleles of *m* loci from *G*_1_ and then mixing with alleles of the other *m* loci from *G*_2_. Hence if *m < k*_1_ or *m < k*_2_, then there is no chance that the incompatibility *ℐ* appears in the hybrid. Otherwise, among all plausible segregation, we can compute the number of achievable ways that the *k*_1_ and *k*_2_ alleles from *G*_1_ and *G*_2_ respectively are sorted into the hybrid. The probability that the hybrid is inviable due to the only incompatibility *ℐ* is thus

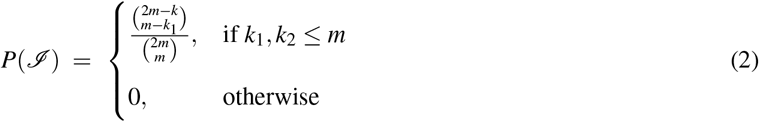

If we further assume that *m* ≫ 1 and *m* ≫ *k*, applying the Stirling’s approximation we have an estimate of the hybrid inviability

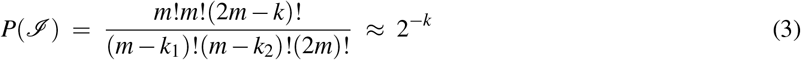

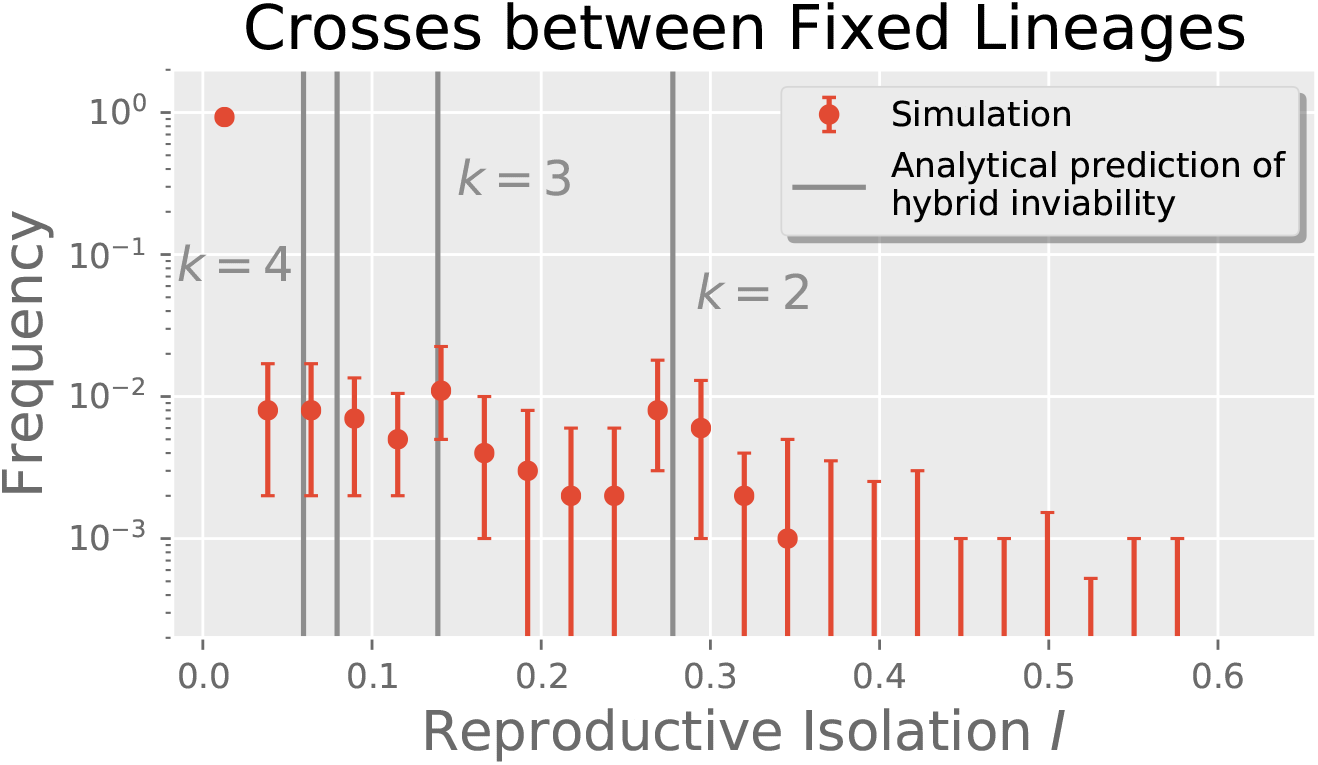
Comparison between the uncovered RI distribution in our simulations and the predicted hybrid inviability Eq 2.

This plain derivation shows that, should there be only one incompatibility concealing between two parental GRNs, the survivability of a hybrid is predominantly determined by the order of the incompatibility.

Here the figure shows good agreement between our analytical prediction of hybrid inviability and the “bulges” from the observed RI distribution. Our simple derivation explains the higher likelihood of certain RI levels relative to their neighboring regions. It also manifests how the discreteness nature of hybrid incompatibilities shapes the RI distribution and that this characteristic has major effects on the strength of reproductive barriers.

## S2 Estimating functional redundancy of GRNs under extreme selection

Our pathway framework not only resonates with existing studies of the functional redundancy of GRNs (True and Haag, 2001; Wagner and Wright, 2007; Schiffman and Ralph, 2018; Láruson et al., 2020), but it also estimates how many GRNs generate a given phenotype under the Boolean-state assumption. Here we consider an extreme case where every protein is either required present or absent, except those that are stimulated by the environment. This scenario depicts a strong selection force, and a weaker selection can be easily reached by relaxing the phenotypic constraint on proteins. Note that this extreme scenario hence provides a lower bound of the number of GRNs that produce the same phenotype.

Suppose there are *n*_+_ and *n_−_* proteins that are required present and absent respectively, and let there be *n*_0_ present state proteins due to the environmental stimuli. A GRN that generates this given phenotype can be viewed as a composition of two parts: First, it contains alleles, i.e., edges, building up pathways from any of the *n*_0_ stimulated proteins to every of the *n*_+_ required-present proteins. Second, edges associated with the required-absent proteins, if any, must not be alleles activated by the required-present/stimulated proteins and producing the required-absent ones. Assuming *m* haploid loci (*m ≥ n*_+_), the number of GRNs generating the given phenotype is

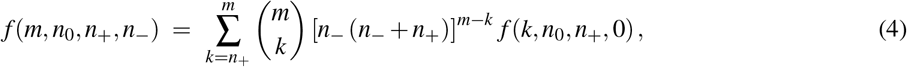

where *f* (*k, n*_0_*, n*_+_, 0) corresponds to the special case where no protein is required absent, it is equivalently the number of directed, edge-labeled graphs with *n*_0_ + *n*_+_ nodes and *k* edges such that every of the *n*_+_ nodes are reachable from any of the *n*_0_ nodes.

Although one may compute compute *f* (*k, n*_0_*, n*_+_, 0) through a recursive relation generalized from existing literature (e.g., Wilf, 2013), an analytical solution is hardly accessible. Here we instead assess the lower and upper bound of *f* (*m, n*_0_*, n*_+_*, n_−_*). First, *f* (*m, n*_0_*, n*_+_*, n_−_*) accounts for all graphs satisfying the reachability criterion, and it is bounded below by the amount of those graphs which are also forests (where a *forest* is a graph which only has trees as its connected components, where trees are graphs without cycles). Finding all such forests is equivalent to finding all possibilities to grow a network from *n*_0_ initial nodes, where edges are added incrementally, pointing from an existing node to a not-yet-existing (newly added) one. So we have

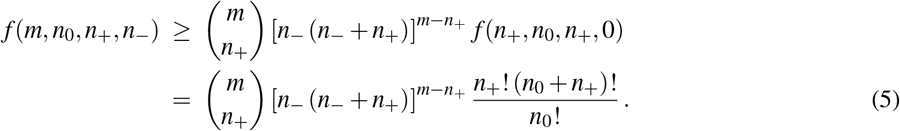

Second, incrementally adding *k − n*_+_ edges to every of such forests is essentially an enumeration of the *k*-edge graphs. This process generates all possible graphs with *k* labeled-edges, but they might be over-counted since adding edges to two different forests can produce the same graph. Computing all possible ways to add *k* edges to every forest satisfying the reachability criterion leads to an upper bound of *f* (*k, n*_0_*, n*_+_, 0):

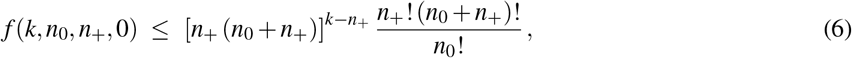

and *f* (*m, n*_0_*, n*_+_*, n_−_*) is hence bounded above by

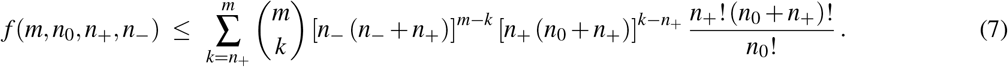

Combined, under an extreme scenario where the binary states of all proteins are constrained, we see super-linearly or even exponentially many GRNs generating the same phenotype. The pathway framework therefore concludes that, for any phenotype derived from the binary states of proteins, the number of functionally redundant GRNs grows faster than super-linearly/exponentially as the system scales.

## S3 Reproductive barriers and ancestral genetic variation

Here we demonstrate our examination on how the extent of ancestral genetic variation influences the appearance and strength of reproductive barriers. To begin with, we designed a pipeline to produce ancestral populations whose amount of genetic variation are tunable. A fixed population was first obtained from our GRN evolution model starting with randomly generated individual GRNs. For every locus, the allele might then mutate into any other possible allele with a per-locus mutation probability *p*. The resulting population was regarded as the ancestral population, where the mutation probability *p* became a tunable parameter to assess the degree of ancestral variation.

We followed the same methodology to simulate generational dynamics of GRNs and to compute reproductive isolation between allopatric lineages as in the main text. Here panel (a-c) of the figure shows, for different number of loci, the reproductive barriers consequent to the varying ancestral mutation probability *p*. Here we present two indicators of barriers: the leading RI (blue, left axis) and the fraction of positive RI (red, right axis). On a first glance the simulations evince that, for a organism with a larger number of loci, the barriers only required a smaller ancestral mutation probability yet more apparent barriers were observed.

Panel (a-c) furthermore suggest some critical level of ancestral variation associated with the constant population size, such that reproductive barriers would hardly appear between lineages evolving from an ancestral population with less polymorphisms. We quantify the critical level of genetic variation through a critical mutation probability *p_c_*; this is the smallest ancestral mutation probability with which a barrier indicator has non-zero median value. Nevertheless, due to the lack of a both sufficient and necessary indicator, we could only estimate the interval that this critical level fell into. The critical level of ancestral variation would be bounded above by *p_c_* for the leading RI (a sufficient indicator of barriers) and bounded below by one for the fraction of positive RI (a necessary indicator of barriers). Panel (d) presents the interval estimation that the critical ancestral variation fell into for organisms with different number of loci.

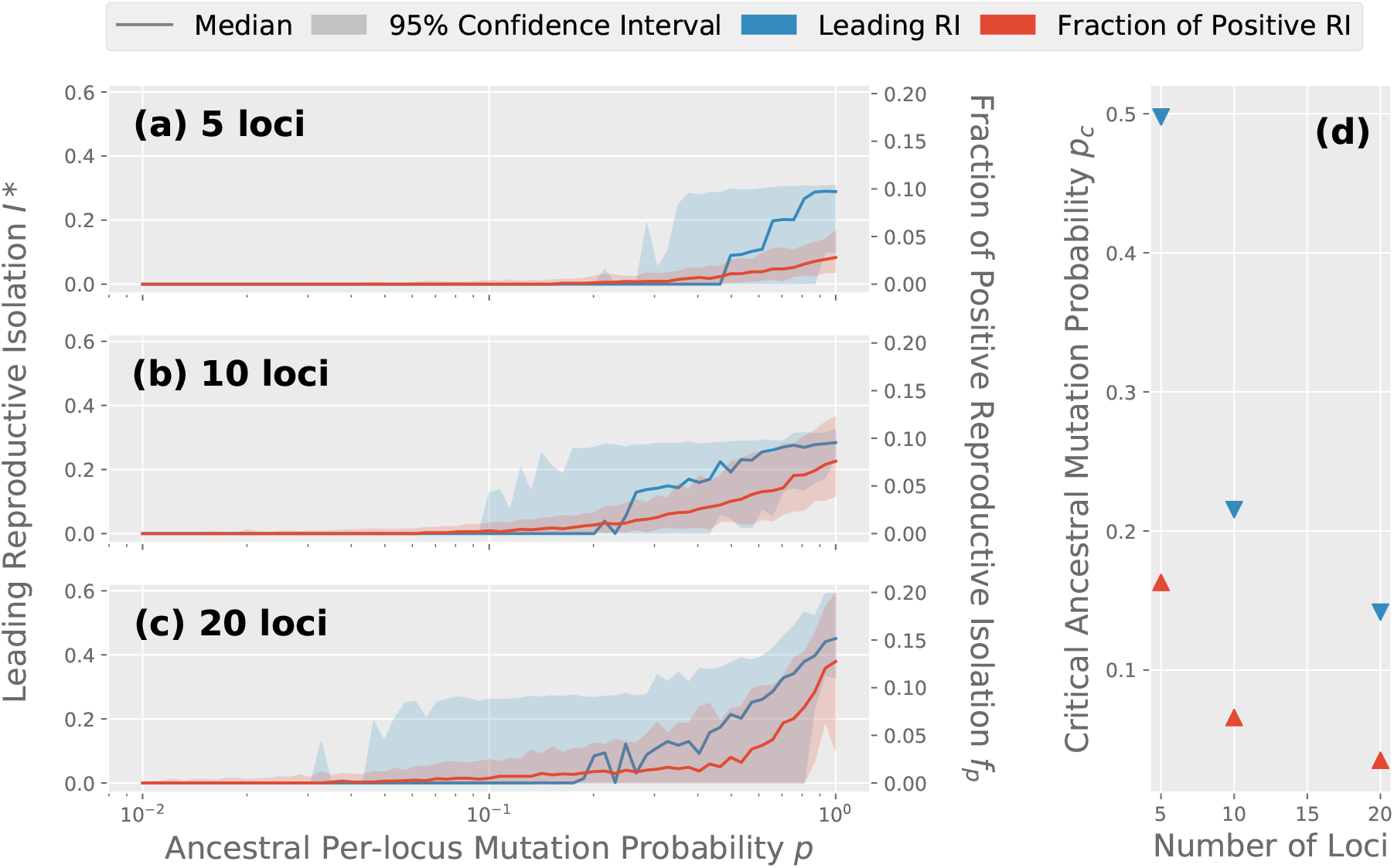
Varying the extent ancestral variation and its corresponding strength of reproductive barriers. The GRN evolution was simulated under Setup 1 described in Methods. **(a-c)** Indicators of barriers for 5, 10 and 20 loci. **(d)** Estimation of their critical level of ancestral variation.

## S4 Algorithms of counting potential incompatibilities

### Algorithm 1 COUNT-ECSP

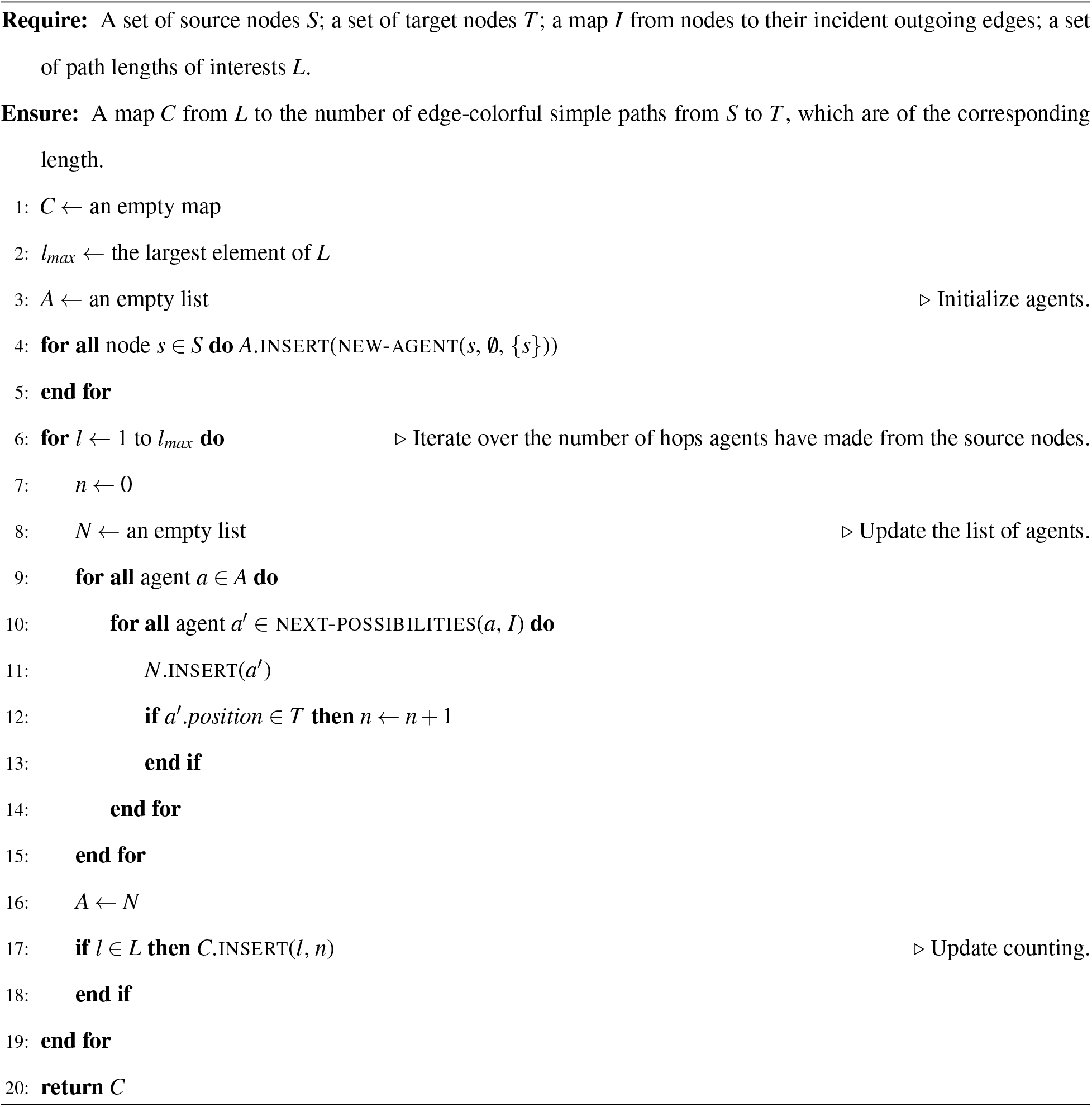

### Algorithm 2 NEXT-POSSIBILITIES

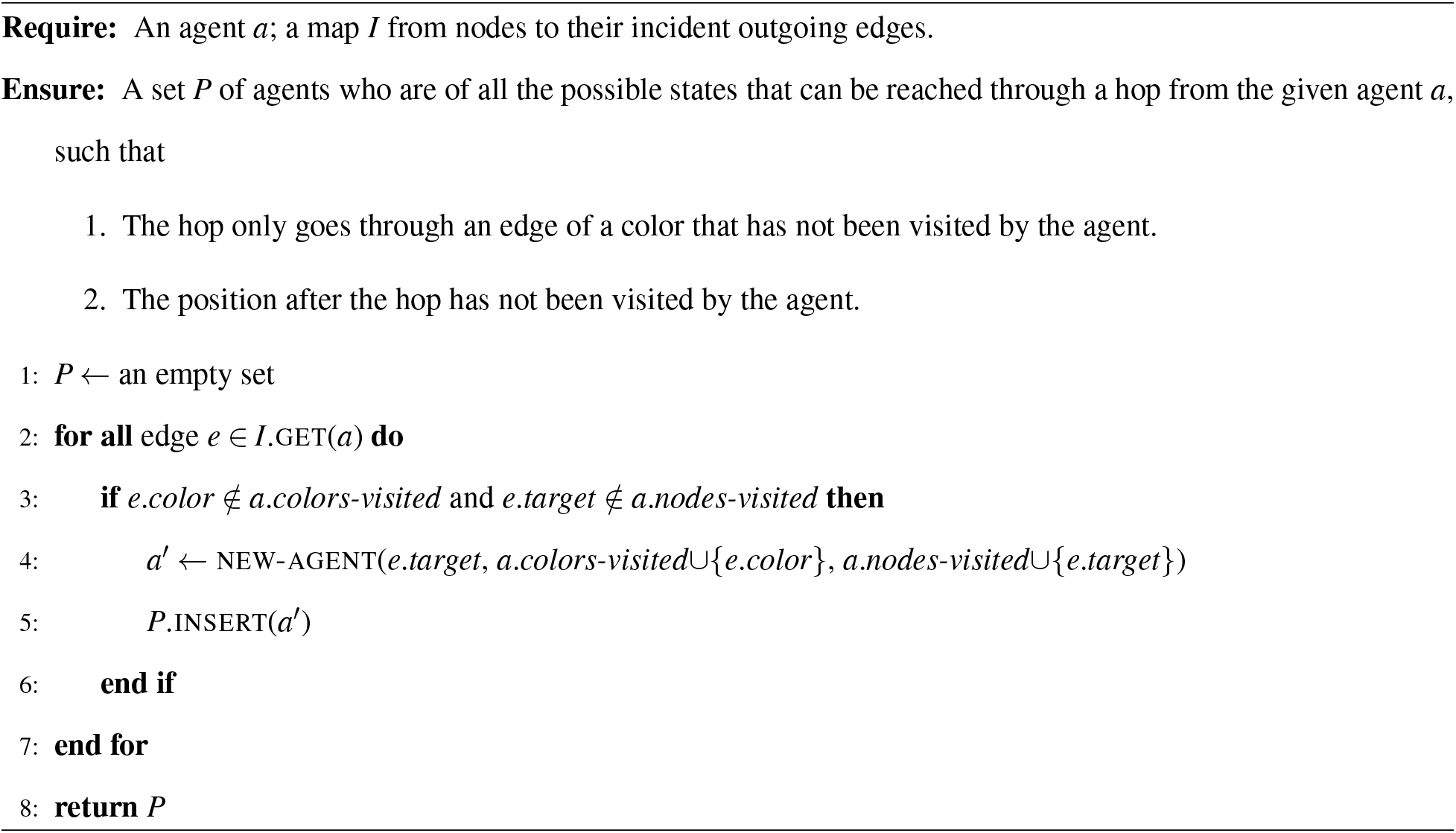

